# TMS of V1 eliminates unconscious processing of chromatic stimuli

**DOI:** 10.1101/606095

**Authors:** Mikko Hurme, Mika Koivisto, Linda Henriksson, Henry Railo

**Author notes:** Corresponding author: Mikko Hurme, Department of Psychology, University of Turku, Finland.

## Abstract

Some of the neurological patients with primary visual cortex (V1) lesions can guide their behavior based on stimuli presented to their blind visual field. One example of this phenomenon is the ability to discriminate colors in the absence of awareness. These so-called patients with blindsight must have a neural pathway that bypasses the V1, explaining their ability to unconsciously process stimuli. To test if similar pathways function in neurologically healthy individuals or if unconscious processing depends on the V1, we disturbed the visibility of a chromatic stimulus with metacontrast masking (Experiment 1) or transcranial magnetic stimulation (TMS) of the V1 (Experiment 2). We measured unconscious processing using the redundant target effect (RTE), which is the speeding up of reaction times in response to dual stimuli compared with one stimulus, when the task is to respond to any number of stimuli. An unconscious chromatic RTE was found when the visibility of the redundant chromatic stimulus was suppressed with a visual mask. When TMS was applied to the V1 to disturb the perception of the redundant chromatic stimulus, the RTE was eliminated. Based on our results and converging evidence from previous studies, we conclude that the unconscious processing of chromatic information depends on the V1 in neurologically healthy participants.

## 1. Introduction

Clinical cases of blindsight suggest a fundamental dissociation between conscious visual perception and unconscious visually guided behavior (Pöppel, Held, & Frost, 1973; Weiskrantz, Warrington, Sanders, & Marshall, 1974). Patients with blindsight have a visual field loss due to primary visual cortex (V1) lesions, but they can process visual information presented to that blind field. Multiple manifestations of these residual capacities are assumed to be caused by different pathways bypassing the V1 (Dankert & Rosetti, 2005), but all these have two aspects in common: V1 lesions and the ability to guide behavior based on visual information that the patients reportedly do not see. This phenomenon has led to the conclusion that unconscious visual processing does not rely on the same neural processes and brain areas as those used in conscious vision. In this study, we investigated whether neurologically healthy individuals could unconsciously guide their behavior based on chromatic information that they do not consciously perceive due to the transcranial magnetic stimulation (TMS) of their V1. Studies on patients with blindsight have revealed that the patients can discriminate colors that they do not consciously perceive (Brent, Kennard, & Ruddock, 1994; Stoerig & Cowey, 1989, 1992) and that unconscious processing of chromatic information can prime faster reaction times (Tamietto et al., 2010). However, it is impossible to generalize the findings from patients with blindsight to the healthy population because new connections are formed and existing connections are modified due to neural plasticity after the lesion (Payne & Lomber, 2001; Leh, Johansen-Berg, & Ptito, 2006). Blindsight may be the result of these neuronal changes rather than a property of an intact human brain.

Using TMS makes it possible to study the causal role of a brain area in a specific task performed by neurologically healthy individuals. TMS induces electrical activation in the stimulated cortical area, which can temporarily disrupt normal processing in the area. The TMS of the V1 can suppress the conscious perception of a stimulus (Amassian et al., 1989), demonstrating the same effect that lesion studies (Holmes, 1918) have shown—the V1 performs a causal role in conscious perception. Since then, studies have tried to demonstrate that after suppressing the conscious perception of a stimulus with TMS pulses to the V1, it is possible to reveal “TMS-induced blindsight” in neurologically healthy observers. Some of the studies have found evidence of this phenomenon with varying stimulus types (Koenig & Ro, 2018; Boyer, Harrison, & Ro, 2005; Railo & Koivisto, 2012; Ro, Shelton, Lee, & Chang, 2004; Christiansen, Kristiansen, Rowe, & Nielsen, 2008), but a larger number of the studies have concluded that both conscious and unconscious visual processing rely on the V1 in neurologically healthy individuals (de Graaf, Koivisto, Jacobs, & Sack, 2014; Hurme, Koivisto, Revonsuo, & Railo, 2017, 2019; Koivisto, Henriksson, Revonsuo, & Railo, 2012; Koivisto, Mäntylä, & Silvanto, 2010; Persuh & Ro, 2013, Railo, Andersson, Kaasinen, Laine, Koivisto, 2014; Sack, van der Mark, Schuhmann, Swarzbach, & Goebel, 2009)

In this study, we employed a method called the redundant target effect (RTE) to measure the effects of unconscious stimuli on behavior. The RTE involves speeding up reaction times when a participant is presented with two stimuli instead of one in a paradigm where the task is to respond as fast as possible when a single stimulus or two stimuli are presented. During the two-stimulus condition, the additional target is called the redundant target because it is not necessary for the task performance. The RTE has been found even when the redundant target is reported as unconscious in patients with V1 lesions (Marzi, Tassinari, Aglioti, & Lutzemberger, 1986; Tamietto et al., 2010; Tomaiuolo, Ptito, Marzi, Paus, & Ptito, 1997) and in neurologically healthy participants (Hurme et al. 2017, 2019; Savazzi & Marzi, 2002). A promising candidate that can explain the unconscious RTE is a neural pathway that connects the superior colliculus (SC) to the extrastriate cortex and bypasses the V1 (Savazzi & Marzi, 2004). The role of this pathway in unconscious processing is very difficult to verify in humans. Sumner, Adamjee, and Mollon (2002) propose that S-cone activating short-wavelength color stimuli can be used to test the contribution of the SC because studies on primates have suggested that the SC is not activated by short-wavelength stimuli (Marocco & Li, 1977; Schiller & Malpeli, 1977; de Monasterio, 1978).

This short- versus long-wavelength paradigm has been used to study the role of the SC in the unconscious RTE. As measured with the RTE, blindsight has been found using long-wavelength stimuli, but the RTE is absent in response to short-wavelength purple stimuli (Tamietto et al., 2010). Importantly, Tamietto et al. (2010) have not observed functional magnetic resonance imaging (fMRI) activation in the SC with short-wavelength purple stimuli, but with gray stimuli, SC activation has been observed. The authors conclude that the unconscious RTE without the V1 is possible with long-wavelength stimuli but not with short-wavelength stimuli because the unconscious RTE is mediated by tracts connecting the SC to extrastriate areas. Leh, Mullen, and Ptito (2006) have previously demonstrated that patients with blindsight show the RTE of achromatic stimuli but not of S-cone activating short-wavelength stimuli, but Tamietto et al. (2010) have been the first to show the dissociation between short- and long-wavelength colors in blindsight. The unconscious processing of luminance-masked long-wavelength stimuli found by Tamietto et al. (2010) seems to rule out the possibility that the colliculus can only mediate the unconscious processing of information related to achromatic stimuli. While the SC is not assumed to rely on color-opponent information transmitted to the cortex (White, Boehnke, Marino, Itti, & Munoz, 2009), it is possible that collicular neurons still show some sensitivity to a stimulus wavelength, similar to magnocellular neurons (Lee & Sun, 2009; Chatterjee & Callaway, 2002).

Although Tamietto et al. (2010) have found an interesting dissociation between short and long wavelengths, the conclusion that their results reveal the RTE as mediated by the SC can be questioned. The possibility to eliminate SC activity using short-wavelength stimuli has been recently challenged because express saccades (fast saccades triggered by SC neurons) can be elicited by S-cone activating stimuli (Hall & Colby, 2016), and S-cone isolating stimuli can in fact activate the SC (Hall & Cloby, 2014). Another blindsight case shows an RTE produced by an unconscious achromatic redundant stimulus, but the RTE is absent for both unconscious S-cone activating stimuli and L- and M-cone activating stimuli (Marzi, Mancini, Metitieri, & Savazzi, 2009). These results indicate that blindsight in general is not a systematic finding and that there are individual differences among neurological patients with V1 lesions (Marzi et al., 1986). Furthermore, there is evidence that neurological patients who show blindsight (as measured with the RTE) have functional connections from their V1 lesion-sided SC to ipsi- and contralateral extrastriatal areas, whereas patients with no blindsight ability lack these connections (Leh et al., 2006).

The findings of Tamietto et al. (2010) have not been replicated in neurologically healthy participants (Railo et al., 2014). The TMS of the V1 has been used to disturb the conscious perception of the color stimulus, and the unconscious RTE has been compared between blue and red colors. No RTE has been found with either color when the redundant stimulus is suppressed by TMS. However, Railo et al. (2014) have actually not demonstrated that the stimuli they have used could produce an unconscious RTE even when the V1 activity is not suppressed with TMS. Thus, their finding could merely show that the behavioral paradigm is not sufficiently sensitive to measure unconscious chromatic processing. Similar to the study on patients with blindsight (Tamietto et al., 2010), Railo et al. (2014) have used rapidly flickering luminance masking to ensure that the participants could only rely on chromatic information in the RTE task. The flickering mask, which is a strong, attention-capturing stimulus, might have interfered with the participants’ ability to unconsciously process the chromatic targets. In patients with blindsight, luminance masking may not pose a similar problem because they do not consciously perceive the luminance mask in their blind field. It is impossible to produce long-lasting, unconscious luminance masking with TMS because of the limited strength of TMS suppression. Consequently, the participants consciously perceive the luminance mask, but the color stimulus in the same visual field is suppressed from their consciousness by TMS. With this in mind, we modified the paradigm in the present study to make the luminance mask unnecessary. In contrast to the traditional RTE paradigm, where either one or two stimuli are presented, in the present study, the participants were always presented with two luminance-matched stimuli, but they were instructed to respond only if they saw a chromatic stimulus. We previously used a similar type of paradigm to examine unconscious processing of motion with the RTE (Hurme et al., 2019).

In Experiment 1, we used a metacontrast mask to disturb the perception of the color of the redundant stimulus, and we tested if this unconscious chromatic information still influenced the reaction times. We expected to find unconscious processing using both S-cone activating short-wavelength blue stimuli and M- and L-cone activating long-wavelength red stimuli because unconscious processing of a metacontrast masked color was previously demonstrated (Breitmeyer, Ogmen, & Chen, 2004; Breitmeyer, Ro, Öğmen, & Todd, 2007). In Experiment 2, the behavioral task was identical to that in Experiment 1, but we disturbed the conscious perception of the redundant stimulus’ color by the TMS of the V1. We formulated two hypotheses. First, if unconscious processing of chromatic information does not depend on the V1, we expect to observe an RTE even when the participants do not report perceiving the redundant color target. Furthermore, if the mechanism suggested by Tamietto et al. (2010) is generalized to healthy participants, we expect to observe an unconscious RTE only in response to red (but not blue) stimuli. This dissociation could reflect the contribution of the SC (Sumner et al., 2002; Tamietto et al., 2010) but could also be caused by magnocellular pathways that are largely insensitive to S-cone stimuli (Chatterjee & Callaway, 2002; Lee & Sun, 2009). Second, if unconscious processing of chromatic stimuli depends on the V1, we should find no unconscious chromatic RTE. This hypothesis is consistent with the brain imaging evidence that clearly shows the activation of the V1 during color perception (Schluppeck & Engel, 2002). About half of the cells in the V1 are selective to color; for example, early processing related to color constancy already occurs in the V1 (Gegenfurtner & Kiper, 2003). Unconscious chromatic processing may rely on the V1, similar to conscious processing.

## 2. Methods

### 2.1. Participants

Twenty participants (6 males, mean age = 23.3, SD = 3.6) took part in Experiment 1, where visual consciousness was manipulated using metacontrast masking. The subjects were university students who participated in the experiment to obtain course credits in an introductory psychology course. Eighteen participants (2 males, mean age = 25.5, SD = 4.9) took part in Experiment 2, where visual consciousness was suppressed with the TMS of the V1. The study was conducted in accordance with the Declaration of Helsinki and with each participant’s understanding and written consent. It was also approved by the ethics committee of the Hospital District of Southwest Finland.

### 2.2. Stimuli

The stimuli were presented on a 24” VIEWPixx/EEG LCD monitor with a refresh rate of 120 Hz and a resolution of 1920 × 1080 pixels. The monitor was centered on the participant’s eye level, and the participant was seated 150 cm from it. The stimuli were presented 2° from the fixation cross (which was located in the center of the monitor) in the bottom left and right quadrants. The stimuli were 0.17° dots, presented against a light gray background (68.8 cd/m^2^) for one frame (8.3 ms). The stimulus colors were purplish blue (hereafter, blue), red, and gray. The blue stimulus (12.9 cd/m^2^, colorimetric values: x = 0.16, y = 0.07) was the same for all participants, but the intensities of the red (colorimetric values: x = 0.64, y = 0.33) and the gray (colorimetric values: x = 0.30, y = 0.30) stimuli were defined for individual participants so that the perceived luminance was the same between the colors.

This was done using a heterochromatic flicker fusion (HFF) task (Lee, Martin, & Valberg, 1988). The participants were presented with a disk (2°) of alternating blue and red colors at a 60-Hz frame rate. They adjusted the intensity of the red RGB channel (all other channels had zero intensity) using a gamepad. When they could not detect the flickering, the red and the blue disks had the same perceived luminance. After adjusting the red color, they performed the same task for the gray stimulus. This time, one increment using the gamepad changed all the RGB channels one bit so that the stimulus always remained achromatic. The HFF was performed twice, first starting with full intensity (fully red or white) and then starting with zero intensity (fully black). The mean of the two obtained HFF values was selected to represent the red and the gray colors in the experiment. In Experiment 1, the measured luminance of the red stimulus was on average 13.3 cd/m^2^ (SD = 2.4), and that of the gray stimulus was on average 14.5 cd/m^2^ (SD = 3.0). In Experiment 2, the luminance of the red stimulus was on average 14.3 cd/m^2^ (SD = 1.3), and that of the gray stimulus was on average 15.9 cd/m^2^ (SD = 2.2).

### 2.3. Procedure

The experiments started with the HFF task (see Section 2.2. Stimuli), where the participants adjusted the colors of the red and the gray stimuli, as described above. A practice block of 40 trials was performed prior to the actual experiment. The experimental task is summarized in Figure 1. Every trial started with a fixation cross, presented at a random interval between 850 and 1,200 ms. Next, two stimuli were presented to the lower visual field, one to the right and the other to the left. The stimuli could be both chromatic or both achromatic or a combination of one chromatic and one achromatic. After the stimulus presentation, the visibility of one of the stimuli was manipulated using either a visual mask 40 ms after the stimulus onset (Experiment 1) or a TMS pulse 90 ms after the stimulus onset (Experiment 2). In Experiment 1, there were also trials where neither of the stimuli was masked to obtain a baseline for the conscious RTE. The participants were instructed to respond by pressing a gamepad button as fast as they could when they saw a chromatic stimulus or stimuli. The red and the blue stimuli were presented in separate blocks so that the participants knew prior to every block what color they were expected to detect. After the stimulus presentation, they were allowed 1200 ms to respond. During this response period, a blank screen was presented. Next, the participants were asked how many chromatic stimuli they saw (none, one, or two) and how confident they were about their number response (confident, quite confident, uncertain, or guessed). The questions and the possible responses were presented on the screen, and the responses were given by pressing gamepad buttons.

**Figure 1.**
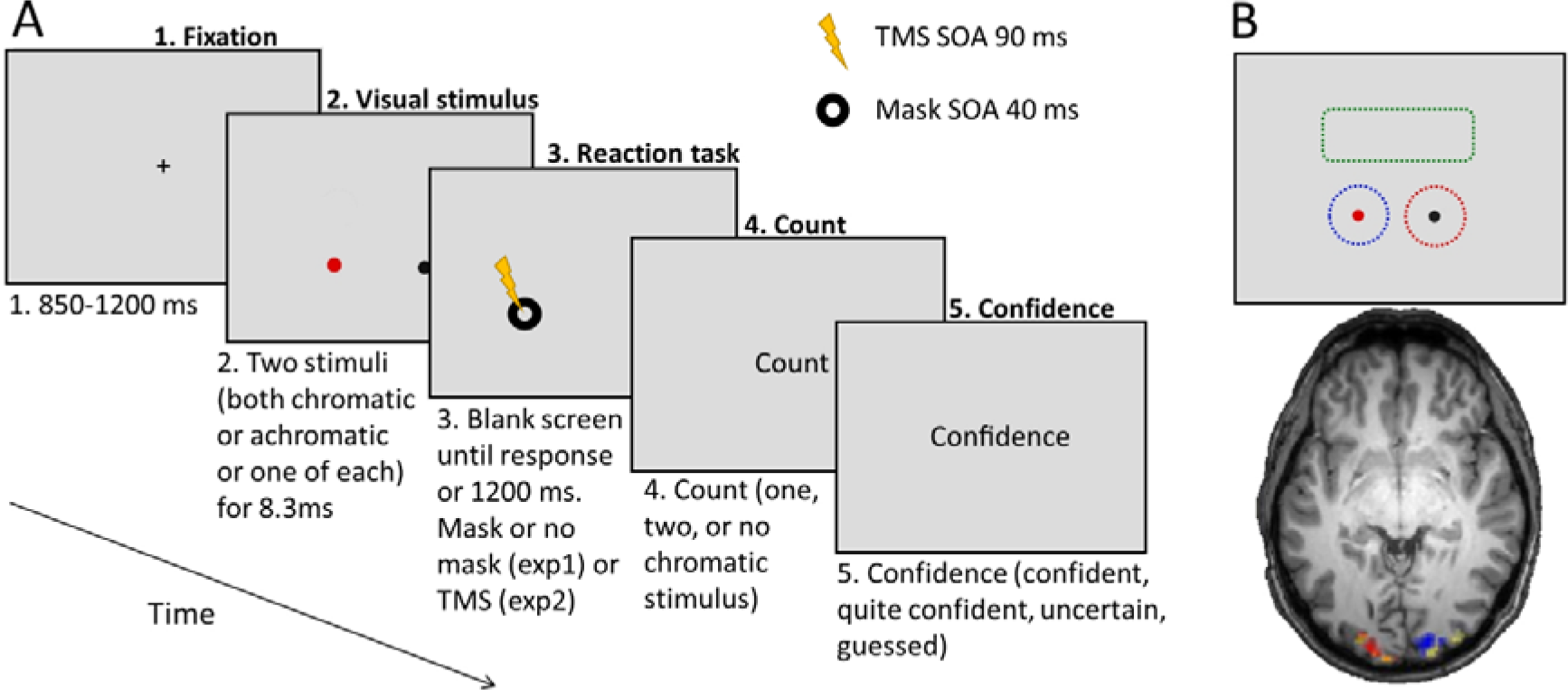
Schematic summary of the experimental procedure. A) Timeline of an experimental trial. The participants’ tasks were to press a button as fast as possible when they detected any number of chromatic targets and then to report the number of chromatic stimuli and their degree of confidence in this decision. The visibility of one stimulus was manipulated using visual masking (Exp1) or transcranial magnetic stimulation (TMS) (Exp2). B) Visual fields (functional magnetic resonance imaging mapped) where processing is disturbed by TMS in Experiment 2. Visual processing in the blue dotted area is disturbed by TMS of the right hemisphere and the processing in the red dotted area is disturbed by TMS of the left hemisphere. The control stimulation target disturbed visual processing in the green dotted area, where no stimuli were presented.

In Experiment 1, the participants completed a total of 24 blocks of 20 trials. They performed the reaction task by first using one hand; after half of the blocks were completed, they changed to the other hand. The red and the blue stimuli were presented in separate blocks, which were done alternately. The starting hand and the order of the stimulus colors were counterbalanced across the participants. In total, 480 trials were completed. In Experiment 2, the participants also finished half of the blocks by using their right hand to perform the reaction task and completed the other half of the blocks by using their left hand. Each hand was used to respond during 6 blocks so that in total, 12 TMS blocks were completed. Each participant had three different stimulation targets in the brain—the left V1, the right V1, and the low center, which served as the control condition (see Section 2.5. TMS). For every stimulation site, the red and the blue stimuli blocks were completed successively for a total of 720 trials. Table 1 presents the number of different trials in the two experiments.

**Table 1.** Number of trials in the experiments. Participants were always presented with two luminance-matched stimuli.

| Experiment | Experiment 1 - Mask |  |  |  |  |  | Experiment 2 - TMS |  |  |  |  |  |
| --- | --- | --- | --- | --- | --- | --- | --- | --- | --- | --- | --- | --- |
| Visual manipulation | No mask |  | Visual masking |  |  |  | Control |  | TMS suppression |  |  |  |
| Target stimulus | Neither |  | Achromatic |  | Chromatic |  | Neither |  | Achromatic |  | Chromatic |  |
| Color (per block) | Blue | Red | Blue | Red | Blue | Red | Blue | Red | Blue | Red | Blue | Red |
| Achromatic stimuli | 48 | 48 | 24 | 24 | - | - | 40 | 40 | 40 | 40 | - | - |
| 1 chromatic stimulus | 24 | 24 | 24 | 24 | 24 | 24 | 40 | 40 | 80 | 80 | 40 | 40 |
| 2 chromatic stimuli | 24 | 24 | - | - | 72 | 72 | 40 | 40 | - | - | 80 | 80 |

### 2.4. Visual masking (Experiment 1)

In Experiment 1, we manipulated the conscious perception of one of the stimuli using a metacontrast mask. The mask was a black annulus whose inner diameter was the same as the outer diameter of the stimulus that it masked. The outer diameter was 0.21° larger than the stimulus. The mask was presented 40 ms after the stimulus onset for nine frames (75 ms). The mask was presented randomly in either the left or the right stimulus. In Experiment 1, there were also baseline trials where no mask was presented. These trials were used to estimate the conscious RTE using this paradigm.

### 2.5. Transcranial magnetic stimulation (TMS, Experiment 2)

TMS was delivered using the MagPro X100 stimulator and the model Cool-B65 coil, a liquid cooled 65-mm figure-of-eight coil, both manufactured by MagVenture. The TMS intensity was set at 85% of the maximum stimulator output. In our previous studies, performed with the Nexstim stimulator, we used a 65–75% stimulation intensity (Hurme et al., 2017, 2019). Pilot experiments suggested that with the MagPro X100 stimulator, approximately 85% intensity is required to reach equally strong visual suppression as with the Nexstim Eximia stimulator. TMS was delivered 90 ms after the stimulus onset. This stimulus onset asynchrony (SOA) was selected because it had been the most effective in suppressing conscious perception in our previous studies, and the classical dip around 100-ms SOA (Amassian et al., 1989) had been shown as the most reliable way to suppress conscious perception with TMS (de Graaf et al., 2014).

The stimulation location was based on the individual participants’ magnetic resonance imaging (MRI) results using the Localite TMSNavigator 3.0.48 system. The targeted stimulation areas within the V1 were defined using retinotopic mapping in fMRI (see Section 2.6. *Magnetic resonance imaging and retinotopic mapping*). For individual participants, the retinotopic locations in their V1, corresponding to the visual field locations where the targets were presented, were the stimulation targets. For all participants, the targets were in the upper bank of the calcarine sulci, in the right hemisphere for the left visual field and vice versa for the right visual field (see Figure 1B). We also included a control target in the visual cortex. The control target was defined anatomically; pulses were delivered to the longitudinal fissure between the lower banks of the calcarine sulci. Consequently, TMS should only affect the upper visual field in the control stimulation, otherwise corresponding to the experimental conditions, where the early visual cortex was stimulated, and the clicking noise and the tactile feedback in the back of the head were similar to the experimental conditions.

### 2.6. Magnetic resonance imaging and retinotopic mapping

MRI was performed in the Turku PET Center using 3T Philips MRI. For each participant, a high-resolution (voxel size = 1 mm^3^) T1-weighted anatomical image of the whole head was captured (3D TFE). Visual cortical areas were mapped with fMRI by using a modified version (Henriksson, Karvonen, Salminen-Vaparanta, Railo, & Vanni, 2012) of the multifocal procedure described by Vanni, Henriksson, and James (2005). The visual stimuli were presented to the scanner using the VisualSystem HD (NordicNeuroLab, Bergen, Norway) binocular apparatus (1920 × 1200 resolution) and the Presentation software (Neurobehavioral Systems, Inc., Albany, CA, USA). For the retinotopic mapping, the major imaging parameters were a 1.8-s repetition time, a 3-ms echo time, a 60°-flip angle, a 25-cm field of view, a 96 × 96 matrix, and a 2.5-mm^3^ voxel size. Twenty-nine slices were acquired in interleaved order. Standard preprocessing with slice-time and motion correction was followed by the estimation of the general linear model with the SPM8 Matlab toolbox.

### 2.7. Data analysis

Behavioral data was analyzed using R statistical software 3.5.0 (R Core Team, 2018); for Bayesian analyses, we used JASP version 0.8.6 (JASP Team 2018, https://jasp-stats.org). Linear mixed-effect models were fitted using the lme4 package (Bates, Mächler, Bolker, & Walker, 2015). The use of mixed-effect models allowed us to take into account the individual differences among the participants as random effects. When selecting the mixed-effect models, we started with the maximum fixed-effect and the random-effect structures. The model with the maximum fixed-effect structure included the following factors: redundant target, stimulus color, and interaction between redundant target and stimulus color. In the second most complex model, the interaction was dropped, and in the simplest model, we only had the redundant target. Random effects were kept maximal as long as the model converged (Barr, Levy, Scheepers, & Tily, 2013). The model with the best Bayesian information criterion (BIC) was selected. In all RTE analyses, the model with the least factors had the lowest BIC, indicating that there were no main effects for the stimulus color or interactions between the stimulus color and the number of stimuli. In the models, we included only those trials where the participants indicated their confidence in their reported numbers. This decision was made to ensure that only the trials where the participants reported fully conscious or unconscious processing of stimuli would be selected in the analyses. The participants who had trouble detecting the stimuli in the baseline condition (i.e., no visual mask in Experiment 1 and TMS of the control site in Experiment 2) in either of the colors (accuracy of less than .8) were removed from the analyses. Based on these criteria, the data from three participants in Experiment 1 and four participants in Experiment 2 were excluded from the analyses. The data and the R-scripts are available at https://osf.io/xtqyg/.

## 3. Results

### 3.1. Experiment 1: Visual masking

#### 3.1.1. Response errors

We instructed the participants to respond only when they saw a chromatic stimulus. Therefore, every trial where only achromatic stimuli were presented and the participant responded was considered a response error. The participants erroneously responded to the achromatic-only trials 8% of the time on average (SD = 6). This low amount of errors indicated that the participants performed the task as intended.

#### 3.1.2. Effects of visual masking

We plotted the accuracies of the number responses and their confidence intervals (Figure 2A) to examine how effective the metacontrast mask was in suppressing the conscious vision. The mask was not very effective in suppressing the conscious perception when only one chromatic stimulus was presented. However, in approximately half of the critical two-stimulus trials, the participants reported being confident that they saw only one chromatic stimulus. We assume that this difference can be explained by the top-down modulation (Ramachandran & Cobb, 1995). When two stimuli are presented and one of them with a metacontrast mask, the unmasked stimulus signal is so strong that it draws attention to it, and the masked stimulus is more likely to be unnoticed. When only one stimulus is presented, the weak signal from the masked stimulus is enough to draw attention to it when no competing stimulus is presented. The mask had little effect on the degree of confidence, as shown in Figure 2B, indicating that visual masking yielded an unequivocal experience of only one stimulus.

**Figure 2.**
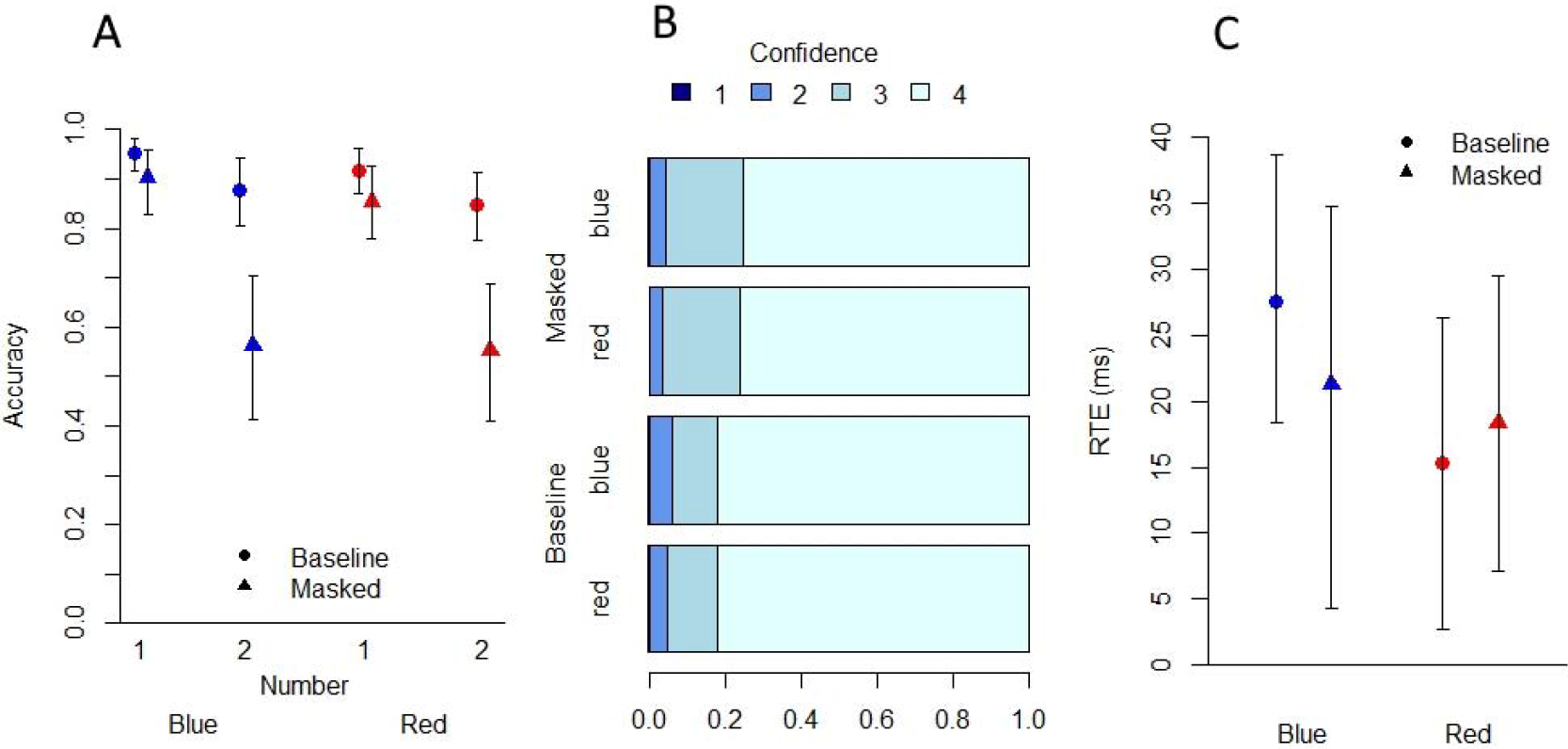
Results of Experiment 1. A) The accuracy of the number response in one-stimulus and two-stimulus conditions with blue or red stimulus colors. B) Proportions of the confidence ratings in baseline and masked conditions for both colors (pooled data from one-stimulus and two-stimulus conditions). C) Redundant target effect in baseline and masked (fully unconscious redundant target) conditions for both colors. The baseline here refers to a condition where no mask is presented. Error bars represent bootstrapped 95% confidence intervals.

#### 3.1.3. Redundant target effect (RTE) during conscious perception of stimuli

Because the RTE task used in this study differed from the typical RTE paradigm, we first wanted to demonstrate that our paradigm would produce an RTE when the stimuli were consciously perceived. The RTE that was calculated based on aggregated data (each participant’s mean RTE) can be found in Figure 2C. The final model with the best BIC included fixed and random effects for the redundant target. As shown in Table 2, the consciously perceived redundant target speeded up the reaction times by 23 ms on average (by participant SD = 9 ms, as indicated by the corresponding random effect).

**Table 2.** Results of the linear mixed-effect model for the conscious RTE baseline

|  | Fixed effects |  |  | Random effects |
| --- | --- | --- | --- | --- |
|  | Estimate ms | 95% CI | T-value | SD |
| Intercept | 285.68 | 271.40 - 304.67 | 35.11 | 31.84 |
| Redundant target | -23.45 | -26.77 - -8.54 | -5.53 | 8.89 |

#### 3.1.4. RTE with an unconscious redundant target (visual masking)

We wanted to find out if the participants could unconsciously process the redundant stimulus, whose perception was disturbed by visual masking. We compared the reaction times between the confidently and correctly perceived one chromatic-stimulus trials and the reaction times in the trials where two chromatic stimuli were presented, but the participants reported being confident (highest confidence rating) that they saw only one. This left us with 82 (SD = 42) trials per participant on average (on average, 19 with one blue stimulus, 22 with two blue stimuli, 18 with one red stimulus, and 23 with two red stimuli). Importantly, in the one-stimulus condition, in addition to the chromatic target stimulus, a redundant *gray* luminance-matched stimulus was masked. The participants’ subjective experience was therefore the same in both situations, and the only difference between those two conditions was the color (chromatic or achromatic) of the masked redundant stimulus. The aggregated RTEs are presented in Figure 2C. The mixed-effect model with the best BIC included fixed and random effects for the redundant target. The results are presented in Table 3. The unconsciously perceived redundant chromatic target speeded up the reaction times by 18 ms on average.

**Table 3.** Results of the linear mixed-effect model for the unconscious RTE (masked)

|  | Fixed effects |  |  | Random effects |
| --- | --- | --- | --- | --- |
|  | Estimate ms | 95% CI | T-value | SD |
| Intercept | 288.03 | 271.40 - 304.67 | 33.94 | 32.84 |
| Redundant target | -17.65 | -26.77 - -8.54 | -3.80 | 7.61 |

### 3.2. Experiment 2: TMS

#### 3.2.1. Response errors

The participants erroneously responded to the achromatic stimuli only 7% of the time on average (SD = 5), indicating that the participants did the task as instructed.

#### 3.2.2. TMS suppression

We plotted the accuracies of the number responses and their confidence intervals (Figure 3A) to visualize the strength of the TMS suppression of vision. As with visual masking (Experiment 1; Figure 2), the suppressive effect was larger in the two-stimulus trials. Compared with Experiment 1 (Figure 2A), the suppression in the one-stimulus trials was more effective when using TMS than when using the metacontrast mask.

**Figure 3.**
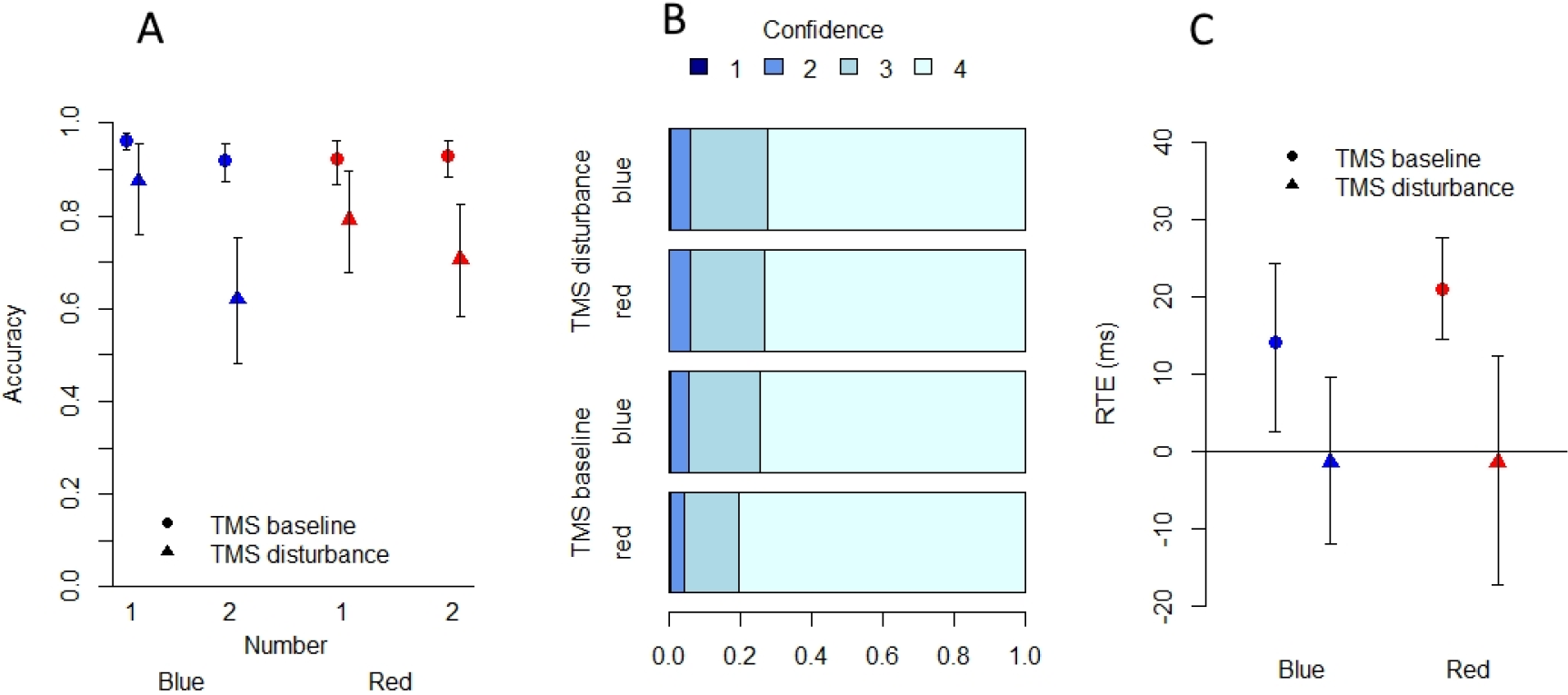
Results of Experiment 2. A) The accuracy of the number response in one-stimulus and two-stimulus conditions with blue or red stimulus colors. B) Proportions of the confidence ratings in TMS baseline and TMS disturbance conditions for both colors (pooled data from one-stimulus and two-stimulus conditions). C) Redundant target effect in TMS baseline and TMS disturbance conditions for both colors. Error bars represent bootstrapped 95% confidence intervals.

#### 3.2.3. RTE baseline under TMS

To obtain a baseline for the RTE in Experiment 2 where TMS was applied, we analyzed the data where the control area in the early visual cortex was stimulated (Figure 1B). We refer to this experimental condition as the TMS baseline. The RTE (calculated from aggregated data, mean per participant) in the baseline TMS condition is presented in Figure 3C, and the distribution of the confidence levels is presented in Figure 3B. In the RTE analyses, we included only those trials where the participants indicated being confident in their reported numbers (highest rating). The final mixed-effect model, selected using the BIC, included fixed and random effects for the redundant target. As shown in Table 4, the consciously perceived redundant target during which TMS was applied to the control area in the visual cortex speeded up the reaction times by 17 ms on average.

**Table 4.**
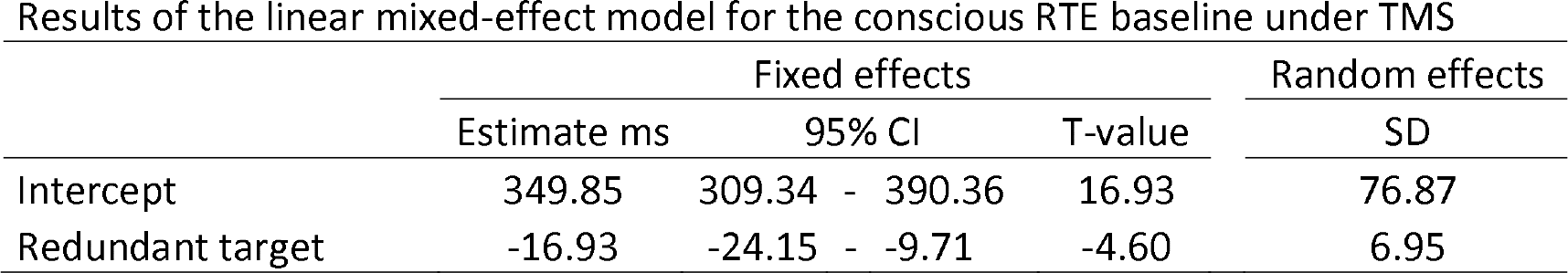
Results of the linear mixed-effect model for the conscious RTE baseline under TMS

#### 3.2.4. RTE when the redundant target is suppressed by TMS of V1

Finally, we investigated whether we could find an unconscious RTE with a color stimulus whose conscious visibility had been suppressed by the TMS of the V1. We call this experimental condition the TMS disturbance condition. The distribution of the confidence in the number response can be found in Figure 3B, and the RTEs (calculated from aggregated data) for both stimulus colors are shown in Figure 3C. In the analysis, only those trials where the participants were confident (highest confidence rating) of their number responses and reported seeing only one chromatic stimulus were included. This left us with 156 (SD = 43) trials per participant on average (on average, 60 with one blue stimulus, 21 with two blue stimuli, 56 with one red stimulus, and 16 with two red stimuli). The best-fitting mixed-effect model was again the one with the least factors, where the response time to a chromatic stimulus was explained by the redundant target. The results are presented in Table 5. When the V1 activity was suppressed using TMS, the redundant chromatic target did not speed up the reaction times. The Bayesian paired sample t-tests for the aggregated means with the uninformative Cauchy prior (scale = 0.707) showed significant evidence for the null hypothesis in both red (BF_10_ = 0.27) and blue (BF_10_ = 0.28) unconscious RTEs.

**Table 5.**
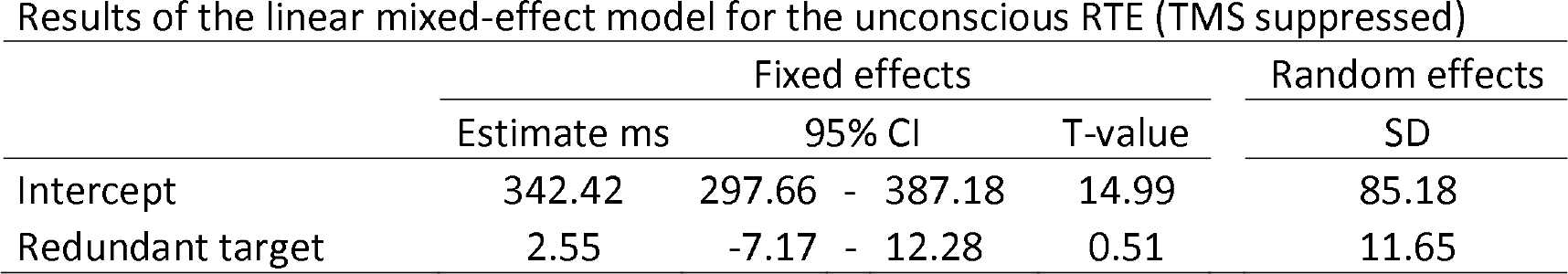
Results of the linear mixed-effect model for the unconscious RTE (TMS suppressed)

|  | Fixed effects |  |  | Random effects |
| --- | --- | --- | --- | --- |
|  | Estimate ms | 95% CI | T-value | SD |
| Intercept | 342.42 | 297.66 - 387.18 | 14.99 | 85.18 |
| Redundant target | 2.55 | -7.17 - 12.28 | 0.51 | 11.65 |

## 4. Discussion

In this study, we demonstrated that unconscious processing of chromatic information from long- and short-wavelength stimuli could influence behavior (Experiment 1), but this would depend on the V1 (Experiment 2). Unconscious processing was measured using the RTE, that is, speeding up the reaction times when two stimuli were presented instead of one. We found an RTE with both colors when the perception of the color of the redundant stimulus was disturbed by a visual mask. However, with the TMS of the V1 to disturb the perception of the color of the redundant stimulus, the unconscious RTE was absent. This result shows that the unconscious chromatic RTE depends on the V1 in neurologically healthy participants. We argue that more generally, converging empirical evidence strongly suggests that the processing of chromatic stimuli depends on the V1 in neurologically healthy humans. Our results are in line with previous studies that have found that unconscious processing of chromatic information depends on the V1 in neurologically healthy individuals when subliminal priming or forced-choice methodology (Railo, Salminen-Vaparanta, Henriksson, Revonsuo, & Koivisto, 2012) or the RTE is used (Railo et al., 2014). Therefore, it seems likely that the unconscious color RTE observed in patients with blindsight (Tamietto et al., 2010) cannot be generalized to neurologically healthy individuals. Another blindsight study has found no unconscious color RTE with short-wavelength or long-wavelength stimuli, indicating that the chromatic RTE is not consistently observed in blindsight, either (Marzi et al., 2009).

It may be argued that the unconscious RTE indexes a highly specific type of unconscious processing; thus, our findings do not rule out the possibility of other kinds of blindsight-like behaviors of neurologically healthy observers. There are two possible explanations for the RTE—neural coactivation and the race model (Miller, 1982)—but both rely heavily on fast responses. According to the race model, the RTE is explained by two signals competing to produce a response that is more efficient compared with one signal. When two signals are presented, the participant responds as soon as either of those signals exceeds the response threshold; therefore, faster responses are more likely. In contrast, the neural coactivation model explains the RTE when the race model is violated. It means that the speeding up cannot be explained by the distribution of the reaction time in response to a single stimuli. The responses to two stimuli are faster than those that could be produced by those two stimuli independent of each other; therefore, it provides evidence for neural coactivation. Either way, the RTE depends on fast automatic responses; it could thus be argued that the unconscious processing of color information without the V1 is too slow to produce the RTE in neurologically healthy participants. However, again, the conclusion is that the unconscious processing of color information by neurologically healthy participants differs from the unconscious processing by patients with blindsight, as reported in the study of Tamietto et al. (2010), who have found blindsight even when using the chromatic RTE paradigm.

What is known about the unconscious processing of chromatic information in paradigms where the TMS of the V1 is used and no speeded-up response is required? An early study using a forced-choice paradigm reports the unconscious processing of color information by neurologically healthy observers when the chromatic stimulus is suppressed by the TMS of the V1 (Boyer et al., 2005). However, there are two methodological shortcomings in this study. First, the study does not apply neuronavigated TMS but a trial-and-error method to find a location in the occipital cortex where TMS suppresses conscious perception. Because the V2/V3 areas are often closer to the scalp than the corresponding visual field locations in the V1, the trial-and-error approach is more likely to stimulate the V2/V3 than the V1 (Salminen-Vaparanta et al., 2012; Thielscher, Reichenbach, Uğurbil, & Uludağ, 2010). Even when TMS targeting is based on retinotopic mapping, the V1 will likely not be *selectively* suppressed, and the V2/V3 representation may also be influenced to some degree (Salminen-Vaparanta et al., 2012). If the geniculostriate visual pathway through the V1 is necessary for unconscious visual processing, then the TMS studies that do not suppress vision by the neuronavigated stimulation of the V1 may observe blindsight-like behavior because they fail to sufficiently disturb the V1 activity. Future studies should examine if blindsight-like behavior is more likely observed when the TMS suppression is produced by the neuronavigated stimulation of the V2/V3 when compared to V1 stimulation.

The second factor that could explain the unconscious processing of chromatic information, as found by Boyer et al. (2005), is their use of a dichotomous seen-unseen consciousness rating that is known to produce false-positive findings. Dichotomous reports are likely to overestimate the amount of unconscious trials (Loyd, Abrahamyan, & Harris, 2013; Mazzi, Bagattini, & Savazzi, 2016; Overgaard, 2011) Railo et al. (2012) measured unconscious processing using a stricter four-step rating scale. Unconscious processing of color information was found when the participants reported seeing something but could not identify the color. However, when they reported being fully unaware of the stimulus (lowest rating), no unconscious processing of color information was found with either the forced-choice or the priming methodology. These results, together with the present findings, suggest that the above-chance response accuracy reported by Boyer et al. (2005) was observed due to the dichotomous awareness rating and was, in fact, a decision based on degraded awareness rather than unconscious perception. In the present study, the participants reported only the number of the *chromatic* stimuli; thus, they possibly retained some residual vision of the stimulus presence (independently of its color). However, in the majority of the trials, the perception of the whole stimulus (not just its color) was most likely suppressed. For instance, in a previous study, we employed a similar paradigm to suppress the visibility of the stimulus completely (Hurme et al., 2017). In the present study, we used a four-point confidence scale so that in the trials where the participants reported being confident about their number answers, they were certain that there were no more or less chromatic stimuli than they indicated. All trials where the participants reported not being confident about their indicated number of chromatic stimuli were excluded from the analyses.

Two TMS studies used the S-cone paradigm (Allen, Sumner, & Chambers, 2014; Railo et al., 2014). Railo et al. (2014) did not find unconscious processing using S-cone stimuli or L/M-cone activating stimuli when they suppressed perception using the TMS of the V1. However, as discussed in the introduction, they failed to demonstrate that unconscious processing of color information was possible with their stimuli when no TMS was applied. Therefore, it might be that their experiment was just insensitive to unconscious processing of color information. Allen et al. (2014) found that they could disturb the conscious perception of both S-cone and achromatic contrast arrow stimuli using the TMS of the early visual cortex, while the accuracy of discriminating the arrow was above the chance level. Based on this result, the authors concluded that they had found the TMS-induced blindsight. However, while the result of the study of Allen et al. (2014) could reflect “relative blindsight”—in the sense that it showed a decrease in subjective visibility without a concurrent decrease in objective performance (Lau & Passingham, 2006)—it differs from the present study in one key aspect. We aimed to use TMS to mimic blindsight in the sense that the V1 activity would be disturbed to the degree that conscious vision would be eliminated. Allen et al. (2014) calibrated the stimuli so that the participants could detect the stimuli only in approximately half of the trials when no TMS was applied. TMS slightly further decreased the visibility of the stimuli, but in most of the trials, the stimuli were not unconscious due to the TMS. Allen et al. had no way of separating the trials that were unconscious due to TMS from those that were unconscious just because the stimuli were weak to begin with. In contrast, we made the task relatively easy in the baseline condition (without TMS) and specifically used TMS to suppress the visibility of the targets.

One limitation of all TMS studies is that the activation that is induced in the brain under the coil spreads to other areas that are connected to the stimulated area (Ilmoniemi et al., 1997). Therefore, it could be argued that the effect of TMS is not because of the disruption in the directly stimulated area but is due to the activations in the connected areas. However, in a similar fashion, local cortical lesions can lead to neuronal changes in distantly connected regions of the brain (Sprague, 1966). Thus, we conclude that the simplest explanation for the present finding is that the suppression of conscious and unconscious vision is due to changes in the V1 activity. We also stress that the control TMS condition was specifically chosen to control for potential network effects. The control stimulation site was the V1, but instead of the upper bank of the V1, we stimulated its lower bank (i.e., V1 but in different receptive field locations). This approach is superior to using a sham stimulation condition (which may feel different to the participant and elicits very weak cortical activations) or a more distant control site, such as the vertex (which feels very different to the participant and activates clearly different networks of brain areas than those in the experimental condition).

A difference between the present study and many studies on patients with blindsight is that we used small stimuli with brief presentation durations. This was done to ensure sufficiently strong suppression by TMS. Studies on patients typically use large stimuli. It thus remains possible that larger and longer-duration stimuli could be more effective in activating neurons in the SC, thereby enabling unconscious chromatic processing. Nonetheless, although the SC has large receptive fields, it is sensitive to very small and brief stimuli (Cynader & Berman, 1972). Wurth, Richmond, and Judge (1980) showed that monkey SC strongly responded to a 5-ms flash (stimulus size = 0.5 × 1.0°). Schiller, Stryker, Cynader, and Berman (1974) demonstrated that a 0.3° flash produced strong activation in monkey SC (stimulus duration = 1 s) when the V1 activity was suppressed by cooling. Furthermore, monkeys with V1 lesions manifest blindsight-like behavior in response to small and briefly presented stimuli (Mohler & Wurtz, 1977), and human patients with blindsight can unconsciously process small and brief stimuli (e.g., Savina & Guitton [2018] used a 0.5° stimulus with an 86-ms duration). We also previously showed that the RTE could be observed when similar small and brief achromatic contrast stimuli were suppressed by the TMS of the V1 at a 90-ms SOA (Hurme et al., 2017). Based on these findings, we suggest that stimulus intensity is likely not the reason why we have not found unconscious chromatic processing, although this question should be directly addressed in future studies.

To conclude, unconscious processing of both short- and long-wavelength chromatic stimuli is possible but not without the V1. This indicates that the neural pathways connecting subcortical structures (e.g., the SC) to the extrastriate cortex, while bypassing the V1, are not sufficient to produce unconscious processing of chromatic stimuli (measured as the RTE) by neurologically healthy participants.

## Acknowledgment

This research was supported by the Academy of Finland (H.R., grant #308533).

